# Efficient Analysis of Proteome-wide FPOP Data by FragPipe

**DOI:** 10.1101/2023.06.01.543263

**Authors:** Carolina Rojas Ramírez, Jessica Arlett Espino, Lisa M. Jones, Daniel A. Polasky, Alexey I. Nesvizhskii

**Affiliations:** Department of Pathology, University of Michigan, Ann Arbor, MI, USA; Department of Pharmaceutical Sciences, University of Maryland, Baltimore, Maryland 21202, USA

## Abstract

Monitoring protein structure before and after perturbations can give insights into the role and function of proteins. Fast photochemical oxidation of proteins (FPOP) coupled with mass spectrometry (MS) allows monitoring of structural rearrangements by exposing proteins to OH radicals that oxidize solvent accessible residues, indicating protein regions undergoing movement. Some of the benefits of FPOP include high throughput and lack of scrambling due to label irreversibility. However, the challenges of processing FPOP data have thus far limited its proteome-scale uses. Here, we present a computational workflow for fast and sensitive analysis of FPOP datasets. Our workflow combines the speed of MSFragger search with a unique hybrid search method to restrict the large search space of FPOP modifications. Together, these features enable more than 10-fold faster FPOP searches that identify 50% more modified peptide spectra than previous methods. We hope this new workflow will increase the accessibility of FPOP to enable more protein structure and function relationships to be explored.

## INTRODUCTION

Protein function arises from three-dimensional structure, making characterizing protein structures and dynamics critical to determining their functions. Mass spectrometry (MS)-based methods are increasingly used to complement traditional structural biology techniques for probing protein structure, as they can provide rapid readouts from small quantities of proteins and tolerate complex mixtures. Protein footprinting methods involve chemical labeling of solvent accessible residues on the protein surface combined with MS-based readout of the modified sites.^1^ One such footprinting method, Hydroxy radical protein footprinting (HRPF) irreversibly modifies 19 of the 20 amino acid side chains. Fast photochemical oxidation of proteins (FPOP)^2^, a form of HRPF using photolysis of hydrogen peroxide to generate hydroxyl radicals, has emerged as a promising method for protein structural characterization^3^. FPOP can be performed even in live cells^4–8^ and animals^9,10^, allowing for perturbations to protein structures to be examined in relevant biological contexts and at proteome-wide scales^11^.

While the ability to irreversibly modify nearly all amino acids is a major advantage of FPOP over other protein footprinting methods, it presents an enormous challenge to interpreting the resulting tandem mass spectra of FPOP-modified peptides. For proteome-scale methods like in-cell and in-vivo FPOP, this challenge represents a significant bottleneck that limits the application of these promising techniques^12^. Proteomics search engines typically consider some common biological and/or chemical modifications, each with a particular amino acid specificity, which are used to generate copies of each base peptide sequence to search for all possible configurations of the modifications on the peptide. The number of possible configurations of a modified peptide grows exponentially with the number of possible modified sites and modifications being considered. When relatively few modifications (and associated amino acid sites) are allowed, the number of configurations remains relatively small, resulting in a manageable search space. FPOP can modify 19 of the 20 amino acids^13^, meaning that the number of allowed modification sites on a peptide is nearly the same as the length of the peptide. Furthermore, many residues have several different modifications that can be induced by FPOP^3,14^, resulting in a total of 51 modification-residue combinations^15^, which further increases the number of possible configurations for a peptide sequence. The result is that search methods that are effective for searching modified peptides in typical proteomics contexts struggle to achieve sufficient speed and sensitivity for FPOP searches. For example, a peptide of length 20 with 1 methionine and 3 serine residues would have 2 possible configurations when considering only methionine oxidation or 16 possible configurations when considering serine phosphorylation and methionine oxidation, but nearly 3.5 billion possible configurations considering an average of 2 possible FPOP modifications per amino acid. Considering all of these possible configurations can be feasible, if time-consuming, using conventional search engines for the digestion of a single protein, but generally fails when expanding to whole-proteome searches that may have millions of digested peptide sequences that each generate billions of possible modified peptide configurations.

Several approaches have been proposed to overcome this enormous search space and enable searches of FPOP-modified peptides. While 19 residues can be modified with multiple possible modifications, in practice, only 14 residues are commonly modified and a most are preferentially observed in a single form^16^. While this still represents a very large search space, conventional search methods have been employed for single/few proteins considering all of these modifications, including Sequest^17^, ProteinProspector^18^, Mascot^19^, PEAKS^20^, and Byonic^21^. In particular, Byonic is notable for allowing modifications to be distinguished between ‘common’ and ‘rare’ forms, allowing more fine-grain control over the number of modifications allowed per peptide. For proteome-wide searches, however, these methods are generally incapable of searching all possible FPOP modifications simultaneously. To avoid this, “multi-node” methods have been developed that split the search into multiple sub-searches, each considering a subset of the possible FPOP modifications, before combining the results to obtain the final identifications^22,23^. This has the effect of disallowing some combinations of modifications to reduce the search space to a manageable size. This method has been successfully applied to whole proteome searches of *C. elegans* in-vivo FPOP data, though it required several hours of search time per raw data file^10^. Finally, search engines that employ fragment-ion indexing^24^, which vastly accelerates conventional searches, have been considered to improve the speed of FPOP searches. The cloud-based search engine Bolt was used to search FPOP data using this approach, however, to fit the modified peptides into computer memory for indexing required several optimizations to the search engine.^15^ While the method primarily used indexed search and a powerful cloud computer to increase the search speed, one of the optimizations employed was to only consider modifications at every 5^th^ amino acid.

We sought to combine an optimized search space with the ultrafast indexed search of our MSFragger search engine to make an FPOP search method that can be accomplished in a matter of minutes per data file. MSFragger offers two ways to search for a peptide modification: “variable modifications” and “mass offsets,” which we combine to create a flexible hybrid FPOP search that offers improved speed and performance. Variable modifications in MSFragger are the conventional method for searching modified peptides, enumerating all possible configurations of the specified modifications for each peptide in the peptide index for fast searching. In contrast, a mass offset search considers only a single modification of a peptide, allowing for many rare modifications to be searched without substantial expansion of the search space (because no combinations of offsets are considered).^25^ Combining these two options allows “tuning” the allowed combinations of modifications in the search: by setting common modifications as variable modifications, multiple common modifications can be allowed per peptide, and by setting rare modifications as mass offsets, a single rare modification can be considered alongside each set of common modifications. FPOP modifications can be swapped between variable modifications and mass offsets depending on their respective frequency in the data. Setting more FPOP modifications as variable modifications increases the number of combinations allowed, increasing the search space and time, whereas setting more as mass offsets reduces the search space and time at the cost of missing some peptides bearing unusual combinations of modifications. We compare a range of settings for this hybrid search to conventional and mass offset searches and to previously published results from a multi-node search of the same dataset. The best hybrid search obtained nearly 259% more PSMs than the multi-node workflow performed in Proteome Discoverer, while also running nearly 57 times faster. Finally, we show that separate score thresholds are needed for appropriate false discovery rate (FDR) control in datasets where FPOP-modified peptides comprise a minority of all spectra. The complete FPOP analysis workflow, including MSFragger search, FDR control, and quantitation, is available in FragPipe 19.1.

## METHODS

### Setup

Raw data from PXD019290 was downloaded from ProteomeXchange^26^ via PRIDE^27^. The dataset is from a study where chemical enhancers were explored to increased hydrogen peroxide uptake in worms for in vivo-FPOP studies.^10^ Briefly, *C. elegans* nematodes were exposed to 200 mM hydrogen peroxide in a homemade flow system. After quenching, nematodes were homogenized followed by ultracentrifugation. The lysate was prepared with typical bottom-up proteomics followed by LC/MS-MS analysis using an Orbitrap Fusion Lumos Tribrid mass spectrometer.^10^ Detailed sample preparation, mass spectrometer parameters, and data analysis can be found in the original publication.^10^ Raw data files were converted to mzML format with MSConvert version 3.0.19296^28^. We analyzed the data with FragPipe (v18.0), MSFragger v3.5, and Philosopher v4.5.1, using Python 3.9.12 for database splitting where necessary. Workflows were run in a Linux server with 28 CPU cores and 2 threads per core. The processor was Intel Xeon CPU E5-2690 v4 @ 2.60GHz and RAM of 500GB. For database searches, a *C. Elegans* database was downloaded from UniProt (ID: UP000001940) with reviewed sequences only (9053 total entries), as well as a second *C. elegans* database containing unreviewed sequences as well (53,404 total entries). Common contaminant proteins and an equal number of reversed sequence decoys were appended using Philosopher.

### MSFragger Searches

MSFragger searches were performed with FPOP-related modifications specified as variable modifications and/or mass offsets (see Table 1 for details). A maximum of 3 variable modifications were allowed per peptide, including FPOP modifications as well as protein N-terminal acetylation. Carbamidomethylation of Cys was set as a fixed modification in all searches. All searches used MSFragger’s built-in mass calibration option, fully enzymatic cleavage with the stricttrypsin enzyme setting (max 2 missed cleavages), peptide lengths of 7-50 amino acids, and N-terminal Met clipping enabled. All mass offset searches used delta mass localization^24^ and reported mass offsets as variable mods in the MSFragger output.

**Table 1.**
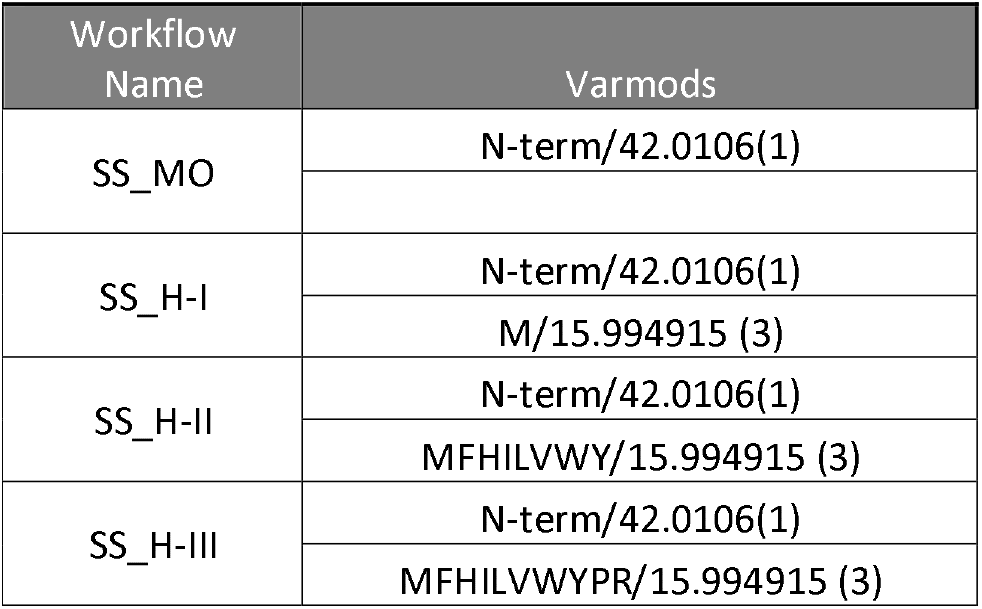
Search space for each FPOP workflow. **All work**flows are identical except in the amount of variable modifications. Workflow SS_MO has only mass offsets and variable modifications are added from workflow SS_H-I to SS_H-III. In the “varmods” column, residues are separated from modification masses by a forward slash and the number of maximum modifications is indicated in parenthesis. For more detailed set of parameters refer to the methods section.

### FDR control for FPOP Database Searches

PSM validation was done using Philosopher (v4.5.1).^29^ FDR calculations were performed using a target-decoy approach with peptide probability calculations first, followed by protein inference and lastly FDR calculations. PSM probabilities were modeled in PeptideProphet^30^ using default closed-search settings (--decoyprobs --ppm --accmass –nonparam –expectscore). Protein inference was performed in ProteinProphet^31^ with default settings. FDR filtering in Philosopher was done to 1% PSM and protein levels, followed by a sequential filtering step which removes any PSMs matched to proteins that are above 1% protein-level FDR. All searches use regular FDR and all FragPipe searches also use group-based FDR filtering to separately filter FPOP-modified and unmodified PSMs. In group-based filtering, prior to sequential filtering, PSMs are separated based into three categories: unmodified, assigned modifications (determined by user input), and other modifications (any modifications other than those specified by the user). Assigned modifications are added in the filter command line as: --**mods M:15.9949**,**n:42.0106**). The benefit of Group-based FDR filtering is to have an appropriate threshold for modified PSMs. In datasets where modified PSMs are not as abundant as unmodified PSMs, performing a regular FDR filtering will set a threshold not appropriate for either PSM category. However, Group-based FDR will set an appropriate threshold for each group.

### Data Analysis

To determined number of PSMs found in each workflow tested, how many PSMs were unmodified, modified, the most abundant modifications and figure making, Python 3.7 scripts were used.

### Data availability

FragPipe output .tsv files obtained for each workflow compare were deposited at https://zenodo.org/record/7823702#.ZGUeAKXMJaR. FragPipe workflow files containing all search parameters used was also provided in the Zenodo Repository. For detailed information in how to understand results tables the reader is directed to https://fragpipe.nesvilab.org/docs/tutorial_fragpipe_outputs.html.

## RESULTS AND DISCUSSION

The benefits of mass offset search have been showcased before in several applications including glycan searches.^24,32^ Increased speed and number of IDs have been observed allowing for new biological insights. In FPOP datasets, the 51 possible residue-modification combinations produce an enormous search space when using conventional variable modification searches. A mass offset search allows only a single modification of a peptide, greatly reducing the search space even if many mass offsets are considered. However, when a peptide is modified with several modifications at once, the expected fragment ions generated by the mass offset search, assuming a modification at a single site, will not match the observed ions in the spectrum (**Figure 1A**, left column). However, setting the less common modifications as mass offsets allows them to be considered, but not more than 1 at a time, reducing the number of combinations to search (**Figure1A**, center column).

**Figure 1.**
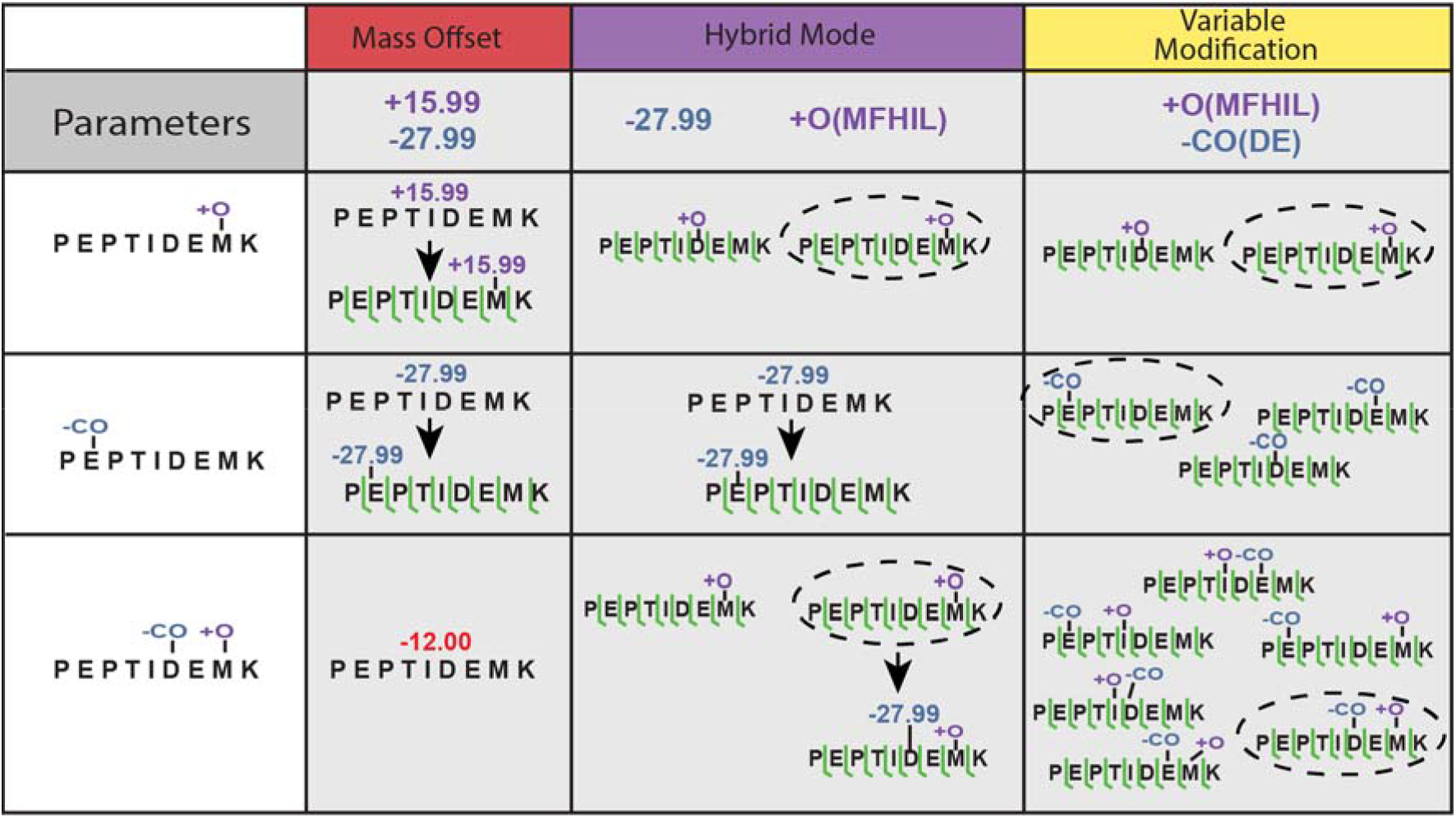
MSFragger hybrid search for improved FPOP PSM identification. Mass offset search considers only a single mass shift/modification per peptide, and thus cannot easily capture multiply modified peptides (left column). By combining mass offsets with variable modifications, some modifications will be captured by the variable modifications and the leftover mass will be quickly identified by mass offsets (center column). The conventional approach, using only variable modifications, can identify multiply modified peptides, but it enumerates all possible combinations of modifications, resulting in very large search spaces (right column).

In the original analysis of Espino et. al, 40,683 FPOP PSMs were identified using a multinode search performed in Proteome Discoverer^10^ (Multisearch with Variable modifications in PD/MS_Var PD). Briefly, the multisearch approach combined results from five individual searches of the data with different modifications considered in each (**Table S1**).^22^ To compare the performance of MSFragger to Proteome Discoverer, we performed this same multisearch approach in FragPipe, using iProphet^33^ to combine the results from each individual search. The FragPipe multisearch (MS_Var FP) identified 91,489 FPOP PSMs, more than twice the PSMs identified by Proteome Discoverer (**Figure 2**). In addition, the MS_Var PD run took more than 96 hours to run, while the MS_Var FP run in FragPipe only took 7 hrs. As a final experiment with conventional search methods, we performed a single node search with all FPOP modifications as variable modifications in FragPipe (SN_Var). This produces a very large search space, as it considers every possible combination of all FPOP modifications at all possible residues. This single node search identified fewer FPOP PSMs than the FragPipe multi-node search and required 168.6 hours, nearly 25-fold longer (**Table S5**). Both the long run time and reduced sensitivity result from the large search space: the long run time intuitively from the large number of modified peptides to check, and the reduced sensitivity because a higher score threshold is needed to separate target and decoy peptides as the search space expands. Therefore, we sought to develop a method capable of efficiently searching FPOP datasets with both fast run times and good sensitivity for a wide adaptability for this information-rich technology.

**Figure 2.**
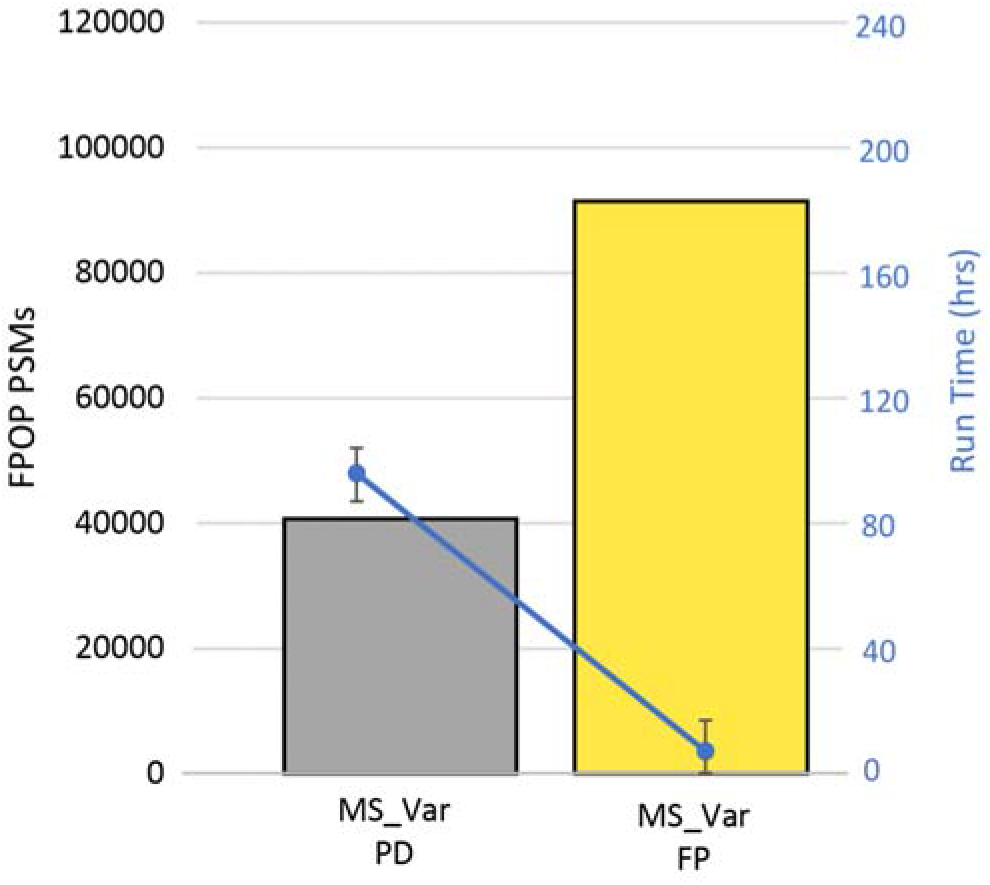
FragPipe produces faster peptide identification workflows with higher performance for FPOP searches. The original dataset was searched using a multinode approach as described previously in Proteome Discoverer (MS_Var/grey), where several FPOP modifications were set as variable modifications.22 Using the same penta level scheme, the same search was performed in FragPipe with double the amount of FPOP PSMs identified (MS_Var/yellow) and 13x less the search time.

Our solution was a “hybrid” variable modification and mass offset search that searches common FPOP modifications as variable modifications (allowing multiple per peptide) and rare FPOP modifications as mass offsets. This restricts the search space by excluding combinations of rare modifications, while still allowing combinations of common modifications to be found. Furthermore, the hybrid search can be “tuned” to the data by switching which modifications are set as variable modifications vs mass offsets, allowing for adaptation to different degrees of FPOP modification in different datasets. To determine the efficacy and speed of these hybrid workflows, we compared three hybrid workflows against a mass offset only search (Single Search with Mass Offset/SS_MO) where all the FPOP modifications were set as mass offsets (**Table 1**). The Single Search with Hybrid search I (SS_H-I) was created by adding methionine oxidation, the most common modification, to the list of variable modifications, and leaving all others as mass offsets. Similarly, workflows SS_H-II and SS_H-III were created by adding oxidation as variable modification for residues FHILVWY and FHILVWYPR, respectively (**Table 1**). For specific mass offsets used in the searches see **Table 2**. From all the hybrid workflows, SS_H-II had the largest number of FPOP PSMs (105,435) identified at 1% FDR, 1.5x more PSMs than MS_Var (**Figure S2A**). For some example MS/MS spectra see **Figure S1**. All the hybrid workflows identified more PSMs than either the multisearch MS_Var or mass offset-only SS_MO searches (**Figure 3A**).

**Table 2.** The FPOP mass offsets used in this study with corresponding modification label and target residues.

| Exact mass | Modification | Residues |
| --- | --- | --- |
| 15.9949 | Oxidation | MFHILVWYADEKNPQR |
| 31.9898 | Dioxidation | CFMWY |
| 47.9847 | Trioxidation | CFWY |
| 13.9793 | Carbonylation | EIKLPQRV |
| -43.0534 | Arg deguanidation | R |
| -22.0320 | H to D | H |
| -23.0160 | H to N | H |
| -10.0320 | H - 10 | H |
| 4.9735 | H + 5 | H |
| -30.0106 | Decarboxylation | DE |
| -27.9949 | CO loss | DE |
| -43.9898 | CO<sub>2</sub> loss | DE |
| -25.0316 | C dioxidation assuming fixed carbamidomethylation | C |
| -9.0367 | C trioxidation assuming fixed carbamidomethylation | C |

**Figure 3.**
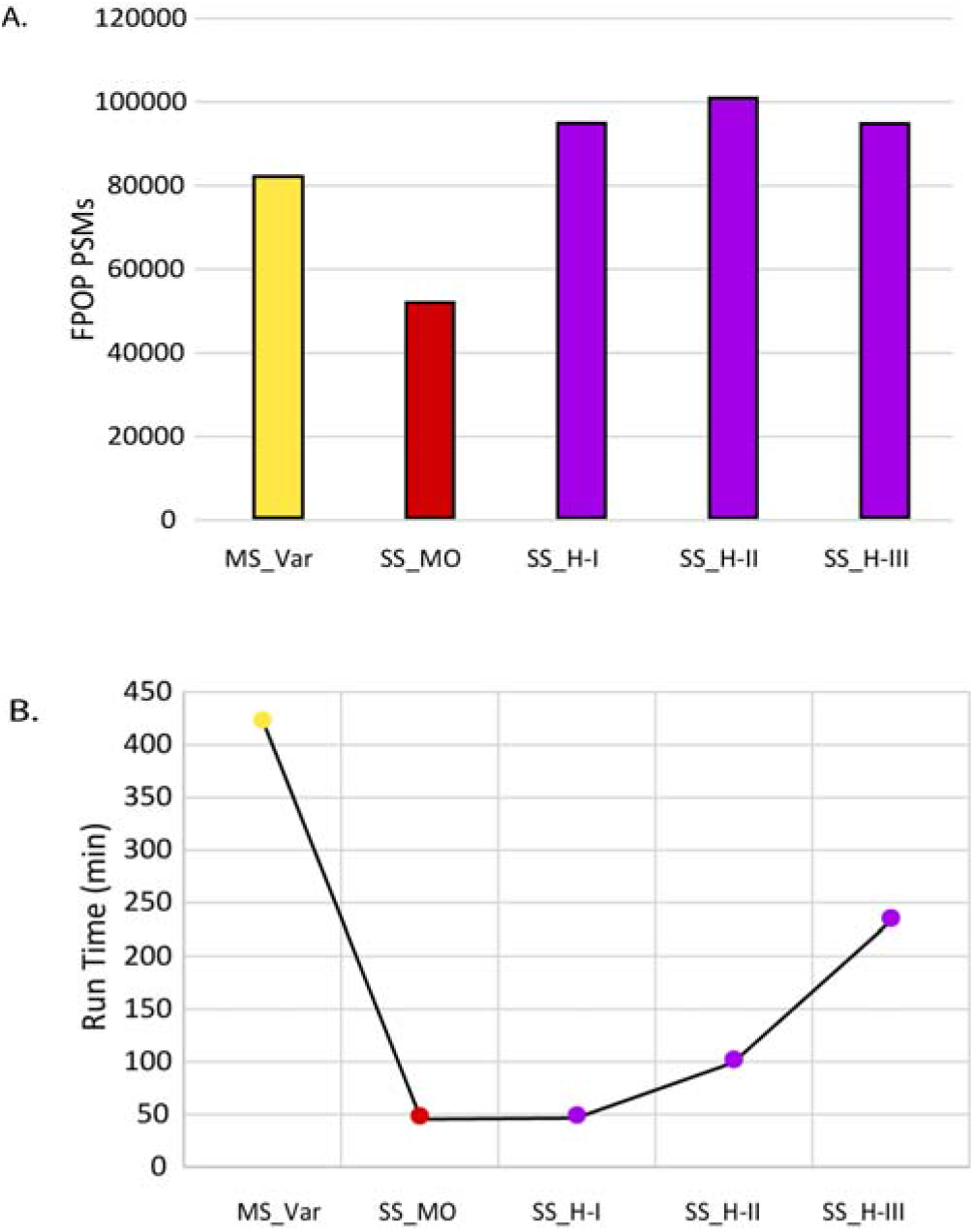
Performance of FragPipe workflows for FPOP datasets. A) The number of FPOP PSMs FPOP PSMs per work-flow using group-based FDR filtering. First the multisearch with only variable modifications (MS_Var) from Figure 2 is shown in yellow. In red, the mass offset only workflow (SS_MO). All the hybrid workflows are shown in purple using the mass offsets shown in Table 2 and the oxidation present in the following residues: (SS_H-I) M, (SS_H-II) MFHILVWY, and (SS_H-III) MFHILVWYPR. FPOP PSMs per workflow using regular FDR filtering are included in the supplementary information section (Table S2). A) Workflow run time. Each workflow is indicated by a label in the x-axis. Each data point corresponding to each workflow is color-coded as A. Run time is in minutes. While SS_MO workflow was the fastest, it identified the lowest amount of FPOP PSMs. All hybrid workflows had faster run times than the multisearch workflow. Run time is shown for regular FDR runs due to being completed continuously.

The large search space for FPOP-modified peptides results in challenges for FDR control methods used in conventional searches. Given the differences in search space for unmodified and FPOP-modified peptides, each requires different PSM score thresholds, as high-scoring decoys (and high-scoring incorrect targets) are increasingly likely as the size of the search space increases. Therefore, we used a group-based FDR filtering method for FPOP data to separately filter unmodified, common modification (defined by the user, in our case, just methionine oxidation), and rare modification-containing PSMs. We chose to include methionine oxidation as a separate category from the other FPOP modifications, as it can occur artifactually during sample prep or from FPOP modification. Given the large background of methionine oxidation in non-FPOP samples, and the fact that it was 26 to 90 times more abundant than the rest of the FPOP modifications (**Table S2**), we separated it from the FPOP modification group to avoid overinflating the confidence of other FPOP-modification PSMs, resulting in a more conservative FDR estimation. However, in datasets with different characteristics, including it with the other FPOP PSMs may be more appropriate. Regular (i.e., all PSMs together) 1% FDR filtering across all four workflows produced an average PeptideProphet probability threshold of 0.79, whereas when separated by group FDR, unmodified PSMs had an average threshold of 0.24, Met oxidation PSMs had an average threshold of 0.16, and the FPOP-modified PSMs had a threshold of 0.98 (**Table S3**). The group FDR filtering reduced the numbers of FPOP-modified PSMs reported in all searches, while increasing the number of unmodified PSMs detected, as expected (**Figure S2B**). Group-based FDR removed the most modified PSMs compared to regular FDR for SS_MO (28%), whereas SS_H-I, SS_H-II and SS_H-III only lost 3, 5 and 7 %, respectively (**Figure S2**). MS_Var lost 11% of FPOP PSMs after group-based FDR filtering. These PSMs were mostly false hits being removed after applying proper FDR thresholds, a further indication that our optimized hybrid workflow produces fewer false hits than mass offset only workflows and conventional FPOP searches.

All workflows were run in the same Linux server as to have the same setup to compare run times. SS_MO, which only used mass offsets for FPOP modifications, had the smallest total run time at only 48 min (**Figure 3B**). As variable modifications were added, the total runtime increased, as expected given the larger search space, reaching a total of 234.8 min for SS_H-III (**Table S4**). The longest workflow, compared to SS_MO, included 10 residues with oxidation as variable modification (**Table 1**). The workflow SS_H-II, with the most PSMs only took 101.5 min, four times faster than MS_Var (**Figure 3B**). Our hybrid method (72 mzML files in less than 2 hrs) is also considerably faster than previous cloud-based high performance search methods for FPOP data (96 mzML files in 64 hrs)^34^.

To demonstrate the capacity to handle large sets of modifications in short amount of time, we performed a search with SS_H-II using a larger database of both reviewed and unreviewed *C. Elegans* sequences, 53,404 total entries^35^. Using workflow SS_H-II to search the unreviewed version of *C. Elegans* identified 288% more FPOP PSMs than using reviewed datasets after group-FDR filtering in 93 more minutes (**Table S7**). Lastly, the addition of N-terminal acetylation did not alter the run time when using SS_H-II to search the unreviewed database and 296% more FPOP PSMs were found compared to SS_H-II (**Figure 4**).

**Figure 4.**
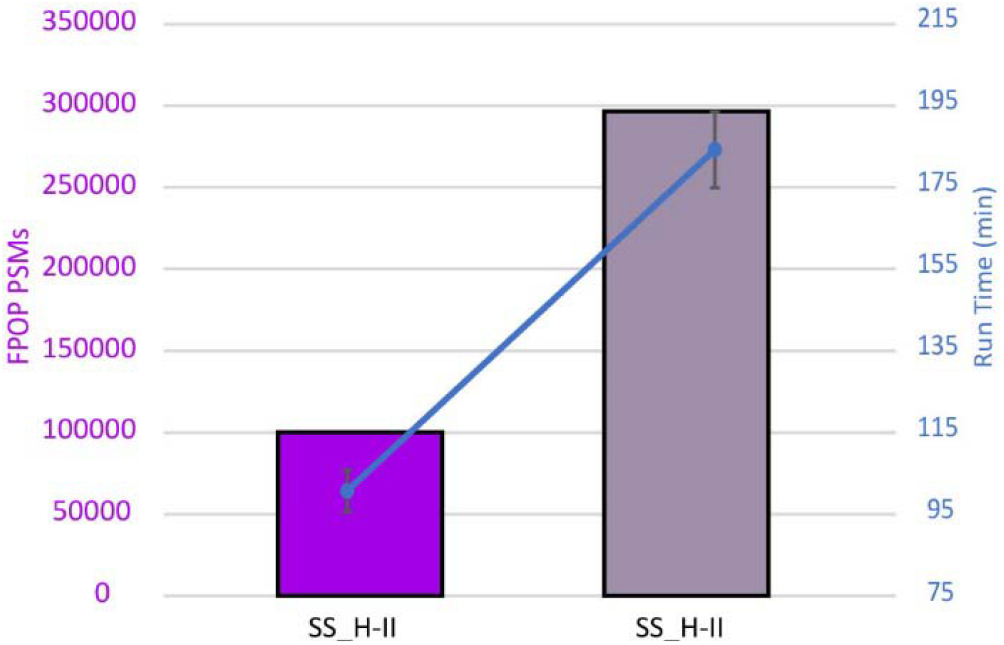
Optimized hybrid FPOP workflow allows for fast searches in larger databases. The number of FPOP PSMs is shown for SS_H-II reviewed (no N-terminal acetylation, reviewed C. Elegans database) and SS_H-II unreviewed+ (N-terminal acetylation, extended C. Elegans database). Superimposed is a graph with the run times for each search showing that the run time is still less than MS_Var FP (7 hrs) even though a database 5.8x larger was used. Data shown is group-based FDR filtered.

## CONCLUSIONS

FPOP has become an important structural biology tool capable of monitoring structural rearrangements in proteins both *in vitro* and *in vivo*. Unfortunately, one of the reasons FPOP lacks widespread adoption is the absence of consensus of an optimal data analysis workflow. Here we present a hybrid search method in FragPipe capable of improving PSM identification at far greater speed. Our hybrid approach contains several variable modifications and sets the rest of the less common FPOP modifications as mass offsets. This ensures that combinations of the most common modifications are identified without expanding the search space to include combinations of rare modifications. Our optimized workflow identified over 100,000 PSMs in approximately 1.5 hours, while the previous Proteome Discoverer workflow identified 40,683 PSMs in more than 96 hours for the same dataset. The optimized FPOP workflow presented here is available in the latest version of FragPipe https://fragpipe.nesvilab.org. (19.1), available at

## ASSOCIATED CONTENT

### Author Contributions

D.A.P. and A.I.N. conceptualization; J.A.E. and L.M.J. raw data production and PD searches; C.R.R, D.A.P. and J.A.E. workflow development; C.R.R and D.A.P. writing: original draft; C.R.R., D.A.P., and A.I.N. writing: review & editing; A.I.N. supervision; A.I.N. funding acquisition.

### Notes

Authors declare no competing financial interest.

## ACKNOWLEDGMENT

This work was funded in part by the National Institutes of Health grants R01-GM-094231 and U24-CA271037.

