## Supporting Information for "Efficient Analysis of Proteome-wide FPOP Data by FragPipe"

**Rapid Analysis of Proteome-wide FPOP Data Using FragPipe.**

^3^Department of Chemistry and Biochemistry, University of California San Diego, San Diego, CA, USA.

| **Table of Contents** | S1 |
| --- | --- |
| **Figure S1.** MS/MS Spectra of PSMs modified with a variable modification, a mass offset, and a combination of both from workflow SS_H-II. | S2 |
| **FigureS2.** Counts for all PSM categories using regular and group FDR Filtering. | S4 |
| **Table S1**. Modifications in multi-node searches. | S4 |
| **Table S2.** Percentage of Methionine oxidation in all modified PSMs. | S4 |
| **Table S3.** Average FDR values and thresholds for all workflows. | S5 |
| **Table S4.** FragPipe Search Times. | S5 |
| **Table S5.** Counts for all PSM categories when using a larger database. | S5 |

**Figure S1. MS/MS Spectra of PSMs modified with a variable modification, a mass offset and a combination of both from workflow SS_H-II.**

A. Variable Modification (M +15.9949)

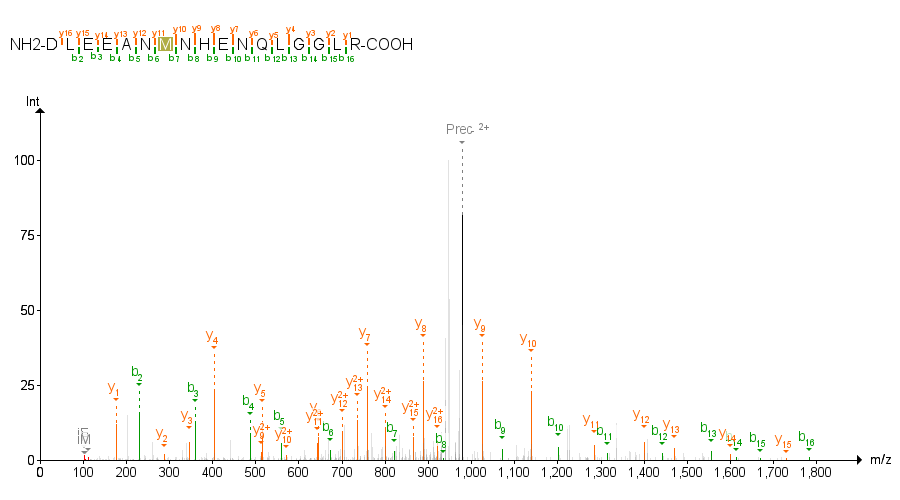

1pAZ_sample_BR2_L2.13395.13395.2_DLEEANMNHENQLGGLR

Assigned Modification: 7M(15.9949)

B. Mass Offset (+13.9792)

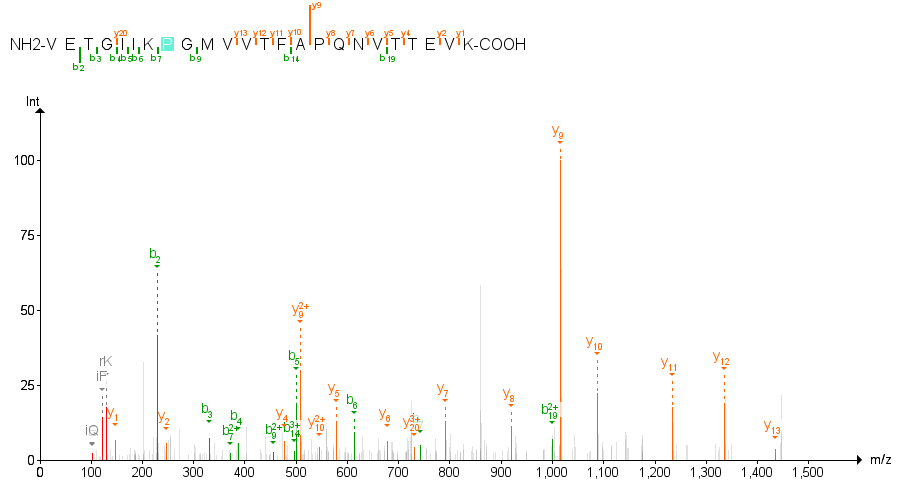

1pAZ_sample_BR1_L2.34085.34085.3_VETGIIKPGMVVTFAPQNVTTEVK

Assigned Modification: 8P(13.9792)

C. Double Modification
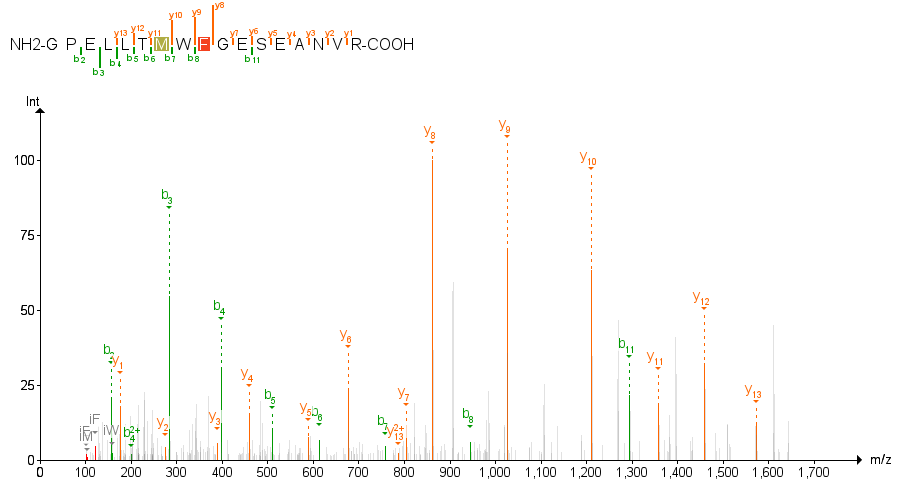

1pAZ_control_BR2_L2.35879.35879.2_GPELLTMWFGESEANVR

Assigned Modification: 7M(15.9949), 9F(15.9949)

**Figure S2**. **PSMs counts A)Regular FDR Filtering.** Due to the low abundance of modified peptides produced in the experimental conditions used, unmodified peptides overwhelmed the signal coming from modified peptides. **B) Group-based FDR filtering.** The PSMs were divided into unmodified group, PSMs modified by methionine oxidation and N-terminal acetylation, and the rest of the modified PSMs.

**A.**

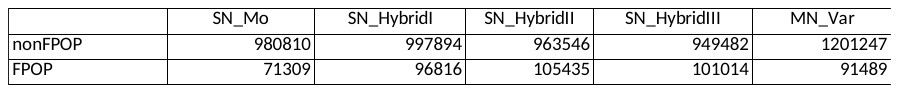

**B.**

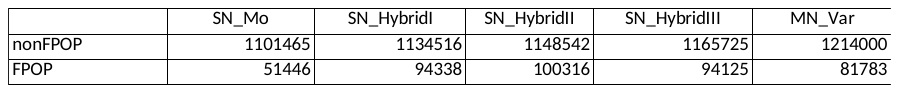

**Table S1. Levels in multilevel search as done previously.**

| node | static mods | variable mods |
| --- | --- | --- |
| 1 | C +57 |  |
| 2 |  | MFHILVWYADENKPQR +16 |
| 3 |  | EIKLPQRV +14 |
|  |  | CFWMY +32 |
|  |  | H -10 |
| 4 |  | CFWY +48 |
|  |  | DE -28 |
|  |  | DE -30 |
|  |  | H +5 |
|  |  | H +5 |
|  |  | R -43 |
| 5 |  | DE -44 |
|  |  | H -22 |
|  |  | H -23 |

**Table S2. Percentage of Methionine oxidation in all modified PSMs.**

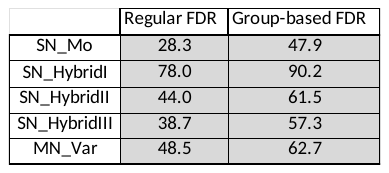

**Table S3.** **Average FDR values and thresholds for all workflows.**

|  | FDR (%) | Threshold |
| --- | --- | --- |
| Regular FDR | 1 | 0.78965 |
| Unmodified FDR | 1 | 0.2419 |
| Defined Modified FDR | 0.935 | 0.16335 |
| All other PSMs FDR | 0.97 | 0.99135 |

**Table S4. Hybrid Workflows Timing.**

| Workflow | SN_Mo | SN_HybridI | SN_HybridII | SN_HybridIII | MN_Var |
| --- | --- | --- | --- | --- | --- |
| Total Time (min) | 48.0 | 48.9 | 101.5 | 234.8 | 420.7 |
| MSFragger (main search only) (min) | 9.9 | 10.7 | 36.9 | 100.1 | 46.9 |
| MSFragger (search and calibration) (min) | 20.0 | 20.6 | 63.5 | 196.1 | 74.8 |
| PSM Validation Step (min) | 28.0 | 28.3 | 38.0 | 38.7 | 345.9 |

**Table S5. Counts for all PSMs categories when using larger databases.** SS_H-II (unreviewed) was done using the unreviewed UNIPROT *C. elegans* database. Unreviewed+ was done with the addition of N-terminal acetylation. The single search, where all FPOP modifications were set as variable (SN_Var) was done with unreviewed database and N-terminal acetylation addition with a run time of approx. 7 days.

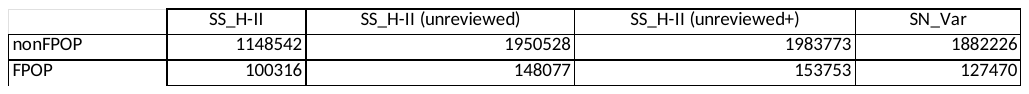
